# The oral cavity of chronically homeless adults is home for unusual extremophile environmental bacteria

**DOI:** 10.1101/2025.04.16.649188

**Authors:** Ali Ruiz-Coronel, Santiago Sandoval-Motta, Daniela Melendez, Violeta Larios-Serrato, Roberto C Torres, Javier Torres

## Abstract

Chronically homeless adults (CHA) often face limited healthcare access, poor nutrition, frequent substance use, and close contact with stray animals. These factors can significantly alter their oral microenvironment. This study aimed to characterize the oral microbiota of this vulnerable and understudied population. We analyzed saliva samples from 60 chronically homeless men and 40 asymptomatic men with no history of homelessness, all living in Mexico City. DNA was extracted, and the V1-V3 regions of the 16S rRNA gene were sequenced. Taxonomic classification was performed using human and non-human databases. Diversity metrics were calculated using the vegan v2.6-4 package in R, and informative species were identified through machine learning and statistical approaches. Microbial correlation networks were inferred using NetCoMi in RStudio. Bacterial diversity was significantly higher in the CHA group. While both groups shared 369 species, only eight were exclusive to the control group, and 149 were unique to CHA. Several of these taxa had never been reported in humans and are typically found in extreme environments. For example, *Megasphaera cerevisiae,* adapted to high ethanol and low pH, was the most abundant species. *Syntrophocurvum alkaliphilum*, from a hypersaline lake in Siberia, and *Sinanaerobacter chloroacetimidivorans*, from anaerobic bioreactors, were also prevalent. Marked differences in microbial network structure were observed between groups. These findings highlight the adaptability of extremophile bacteria to human environments under chronic stress and poor hygiene. This study provides a first look into the oral microbiota of individuals experiencing chronic homelessness, emphasizing the need for more research into marginalized populations.

## Introduction

The oral cavity contains a variety of microenvironments, each supporting specific bacterial communities (Xu et al., 2015). Different regions such as the gingiva, cheeks, palate, tongue, and teeth offer distinct biochemical conditions that shape microbial colonization. For example, surfaces on the tongue (e.g., tonsils, circumvallate papillae, dorsal surface) and various tooth regions host different bacterial profiles. The oral microbiota is among the most diverse in the human body, second only to that of the gut (McLean et al., 2022) (Torres-Morales et al., 2023) (Mark Welch et al., 2020). This diversity is shaped by the oral cavity’s open and dynamic nature, it is continuously exposed to changing nutrients, fluids, gases, temperature, antibiotics, and other external inputs. Host-related factors including genetics, age, diet, lifestyle, and hygiene, also influence the microbial landscape. As a result, the composition of the oral microbiota can vary significantly, even within the same individual over time.

Despite this variability, certain microbial communities are consistently found across individuals. These core oral microbiota are well-adapted to the physicochemical conditions of specific niches in the mouth. Their presence reflects a balance between host factors and environmental exposures that select for stable, niche-specific bacterial clusters.

Among the populations living under extreme environmental and social stress, chronically homeless adults (CHA) offer a unique case to investigate how harsh conditions shape the human oral microbiota. In 2021, the World Economic Forum estimated that 150 million people worldwide experienced homelessness (*Saneamiento*, n.d.), with most living in major cities and concentrated in specific areas. Reliable data are scarce, as this population is often excluded from official statistics and health research. Our ethnographic research in Mexico City revealed that CHA typically lack access to healthcare, bathrooms, clean water, and communication tools like the internet and telephones. Illiteracy is also common within this group and further limits access to services. Diets are poor in quality, based mostly on carbohydrates and fats, with minimal protein.

Oral hygiene is rare, and many individuals consume low-cost, harmful substances daily, increasing their vulnerability to disease. In addition, close and frequent contact with animals (pets) often involving shared food, exposes them to a wide range of microorganisms (*Consultorio abierto “Una puerta de acceso a la salud para personas en situación de calle” - SDSN México*, 2022). These combined factors create a severely disrupted oral microenvironment. To address the lack of healthcare, especially during the COVID-19 pandemic, we launched the “Open Door Clinic” (ODC), a mobile health initiative operating in public spaces where homeless individuals frequently gather. Clinics provided basic medical services and allowed us to document general health conditions. Unexpectedly, we observed very few COVID-19 cases, despite the harsh circumstances. This finding prompted us to investigate the structure and composition of their oral microbiota. Their oral environment, shaped by prolonged exposure to toxic substances, frequent animal contact, shared food, and a lack of hygiene and medical care, represents a highly unconventional microbial niche which to the best of our knowledge, has not been studied before. We believe that characterizing this group may uncover microbial adaptations shaped by prolonged exposure to extreme urban conditions.

## Methods

### Population studied

In this study, individuals were considered chronically homeless adults if they were between 18 and 60 years old and had spent at least three consecutive years living without a stable shelter. (*Website*, n.d.-a). During 2022, the individuals attending the ODCs that met these criteria were invited to participate in the study, and those who agreed provided written informed consent. The final study cohort included 60 CHA and 40 asymptomatic males with no history of homelessness, all residing in Mexico City.

### Clinical data

In the ODC, screening for the detection of COVID-19, information on the disease, primary healthcare, referral to hospital care when needed, and vaccination against COVID-19 and influenza were provided to CHA. Anthropometric measurements, including weight, height, body mass index, and body composition, were recorded. In addition, glucose, blood pressure, oxygen saturation, body temperature, and Human Immunodeficiency Virus and Syphilis tests were offered.

Among the individuals composing the sample, the average age was 51 years, with a minimum of 23 years and a maximum of 80 years. The average weight was 63 kg, with a minimum of 42 and a maximum of 105 kg. The average height was 1.64 m, with a 1.51 to 1.86 m interval. The average BMI was 23.3 kg/m² with an interval of 16.2 to 34.4 kg/m². We found one case positive for syphilis and four positive for COVID-19 using the antibody test. No positive cases were found using the antigen test.

Social and demographic information was also collected from participants, including living conditions, experiences in accessing healthcare services, health literacy, and the challenges that they were facing during the pandemic. The project was approved by the Ethics Committee of the Institute for Social Research at Universidad Nacional Autónoma de México, UNAM, Mexico City.

### Saliva samples

Volunteers were requested to provide a sample of saliva. They were given 10 mL of saline solution (0.85% NaCl), asked to thoroughly wash the mouth, and spit back into a 50 mL plastic tube. Samples were immediately put into ice and promptly transported to a central laboratory for storage at -70° C until tested.

### Preparation of the amplicon libraries from the 16S rRNA hypervariable regions V1-V3 and sequencing

One mL of saliva solution was centrifuged, and bacterial pellets were incubated with lysozyme solution (20 mg/mL, 20 Mm Tris-HCl, pH 8.0; 2 mM EDTA; Triton 1.2%). Next, DNA was extracted using the QIAamp DNA mini kit (Qiagen) following the manufacturer’s instructions. The amplicons of the V1-V3 hypervariable regions of the 16S rRNA gene were generated using previously published primers (Muyzer et al., 1993) ligated to the adapter sequences (Kozich et al., 2013) (Table S2) and used to assemble the DNA libraries. Libraries were amplified using the following conditions: 3 min at 98° followed by 25 cycles (20 s at 98°C, 30 s at 65°C, 30 s at 70°C), 5 min at 72°C and 4°C hold and sequencing was performed on the MiSeq platform (Illumina, San Diego California, USA).

### Bioinformatics analysis

### Raw data processing and normalization

Paired-end sequencing reads in FASTQ format were QC inspected for Phred values and for the absence of adapters using FastQC v0.11.9 (*Babraham Bioinformatics - FastQC A Quality Control Tool for High Throughput Sequence Data*, n.d.). The data were processed using the DADA2 v1.20.0 pipeline (Callahan et al., 2016) with standard filter parameters (maxN = 0, truncQ = 8, and maxEE = 2, reads shorter than 200 bp were discarded). Dereplication filtering and removal of chimera formation was done using the removeBimeraDenovo option. 16S sequences associated with chloroplast or mitochondria were removed. The remaining sequences were grouped into Amplicon Sequence Variants (ASVs) with the naive RDP Bayesian classifier of DADA2, and initial taxonomic classification was assigned to the species level using the Genome Taxonomy Database (GTDB) (Parks et al., 2018).

### Taxonomy assignment refinement

Many ASVs remained unassigned using the GTDB database and were further classified using BLAST searches (Zhang et al., 2000). ASV sequences were then aligned using MUSCLE v3.8.31 and hierarchically clustered based on pairwise SNP distances calculated with snp-dists v0.7.0. ASVs were classified according to the assigned taxonomy of closely related ASVs within the same cluster. Species-level abundances were then aggregated, and raw abundance values were scaled between 0 and 1. Rare taxa, defined as species detected in fewer than two samples, were removed. All steps involving clustering, merging, aggregation, filtering, and normalization were executed in R.

### Alpha and Beta diversity estimation

Alpha diversity was evaluated using species richness, Shannon index, Simpson index, and Pielou’s evenness (Shannon index divided by the natural logarithm of species richness) calculated with vegan v2.6-4 in R. Normality for each index was tested using the Shapiro-Wilk test in R. Differences between controls (housed) and cases (CHA) groups were analyzed using t-tests or Wilcoxon test when normality was not met. Statistical significance was set at P ≤ 0.05.

Beta diversity was assessed using Bray-Curtis dissimilarity and Jaccard distance computed with the vegdist function from the vegan v2.6-4 R package. Principal Coordinate Analysis (PCoA) was then performed using the cmdscale function from the stats v4.2 R package. Results were visualized using ggplot2 v3.4.1, including ellipses representing 95% confidence intervals to illustrate within-group variability.

### Microbial Community Composition

Microbial composition was determined by counting species detected in at least two samples. Shared and group-specific species were identified. Group-specific taxa were defined as species present in at least five samples within one group and absent in the other. Differences in the relative abundance of shared species were evaluated using log2 fold-change analysis, with statistical significance assessed via t-test or Wilcoxon according to data distribution with stats v4.2 R package. p-values were adjusted for multiple comparisons using the Benjamini-Hochberg FDR method; adjusted p-values < 0.05 were considered significant. Shared and group specific species, as well as prevalence and mean relative abundance of group-specific taxa, were visualized with ggplot2 v3.4.1 in R.

### Identification of informative species for Control and Case groups

Informative Species that differentiate controls and cases were identified using machine learning and statistical approaches. A Random Forest (RF) classifier was trained on normalized abundance data using the train function and “rf” method from caret v6.0-94 R package. Model performance was evaluated via 10-fold cross-validation using the trainControl function, and feature importance was calculated with the varImp function of caret. In order to optimize the selection of informative species, model accuracy was assessed across different feature importance percentiles and species below the 85th percentile were excluded. Performance curves and rank plots of the most informative species were generated using ggplot2 v3.4.1 in R.

The beta diversity was re-evaluated using the most informative species selected through the above Random Forest analysis. Additionally, a Linear Discriminant Analysis (LDA) was applied to further investigate the discriminatory power of the selected species. Species with correlation coefficients > 0.75 were removed to improve model performance. The LDA model, implemented using the MASS R package, highlights those species that contributed the most to separate groups. Bar plots were generated with the ggplot2 v3.4.1 R package to visualize the absolute LDA coefficients and the normalized abundances of these species.

### Correlation network analysis

The correlation network was constructed using NetCoMi v.1.1.0 in RStudio v.2024.04.2+764 (Peschel et al., 2021). The netConstruct function was used with the Pearson method as the association measure. To reduce the network to a manageable size (sparsification), a cutoff value of 0.3 was applied, and an additional cutoff of 0.3 was added for local false discovery rate correction. Data was normalized with the modified centered log-ratio transformation (“mclr”). The netAnalyze function was implemented using the “cluster_fast_greedy” method. The graphs were generated with the “spring” layout algorithm. Node sizes were adjusted based on betweenness. Estimated associations are shown in red for negative associations and green for positive associations.

### Metabolic pathway

Pathway prediction was performed using KEGG Orthology at three levels (KO) (Kanehisa & Goto, 2000), and Tax4fun2 was used to obtain relative abundances of pathways (Wemheuer et al., 2020). The z-score of relative abundances and clustering analysis were estimated using the pheatmap package (v1.012). The relative abundances of KEGG pathways were analyzed using linear discriminant analysis (LDA) combined with effect size (LEfSe). The microbial package (v.0.0.21) was used with the ldamarker function with normalization method a. Plots were generated with the plotLDA function using the Kruskal-Wallis test and adjusted *p-*values (padj) < 0.05 and LDA scores ≥ 2.

## Results

After excluding samples with unacceptable quality, 90 saliva samples were sequenced, of which 85 passed the Q30 quality value with an average of 300,000 reads per sample and a total yield of 7 Gb. The initial attempt to assign ASVs with the human microbiota database (GTDB database) left numerous ASVs unassigned in the CHA samples, and the search was extended to other more open bases (including data from the environment). As a result, we learned that CHA samples included several bacteria reported in the environment, including the extremophile species described below.

### The composition of bacterial communities markedly differs between CHA and the control group

Bacterial diversity was significantly higher in CHA for all indexes measured (richness, evenness, Shannon, and Simpson) (Figure 1a) already suggesting important differences between groups. The effect of age, weight, height, and BMI on diversity was analyzed (Figure 1b), and of note, diversity increased as weight and BMI increased in CHA microbiota. In contrast, the behavior was the opposite in controls. The composition of the bacterial communities also showed contrasting results between groups. A PCoA analysis clearly suggested a separation between the groups (Figure 2a), which was confirmed when shared species were determined. Both groups shared 369 species, whereas only eight were specific for controls and as many as 149 were specific for CHA (Figure 2b) many of them not reported previously in humans. Among these 149 we determined the species present in at least five of these individuals and found 43 (Figure 2c). Interestingly, *Megasphaera cerevisiae* was detected with high abundance in as many as 24 CHA individuals, whereas *Prevotella conceptionensis* and two other Bacteroidetes were present in over 15 cases (Figure 2c). Species unusual in humans, like *Syntrophocurvum alkaliphilum*, *Streptococcus downei*, *Thermanaerovibrio acidaminivorans*, *Sinanaerobacter chloroacetimidivorans* or *Desulfovibrionales canine*, were present in 5 to 15 individuals.

**Figure 1.**
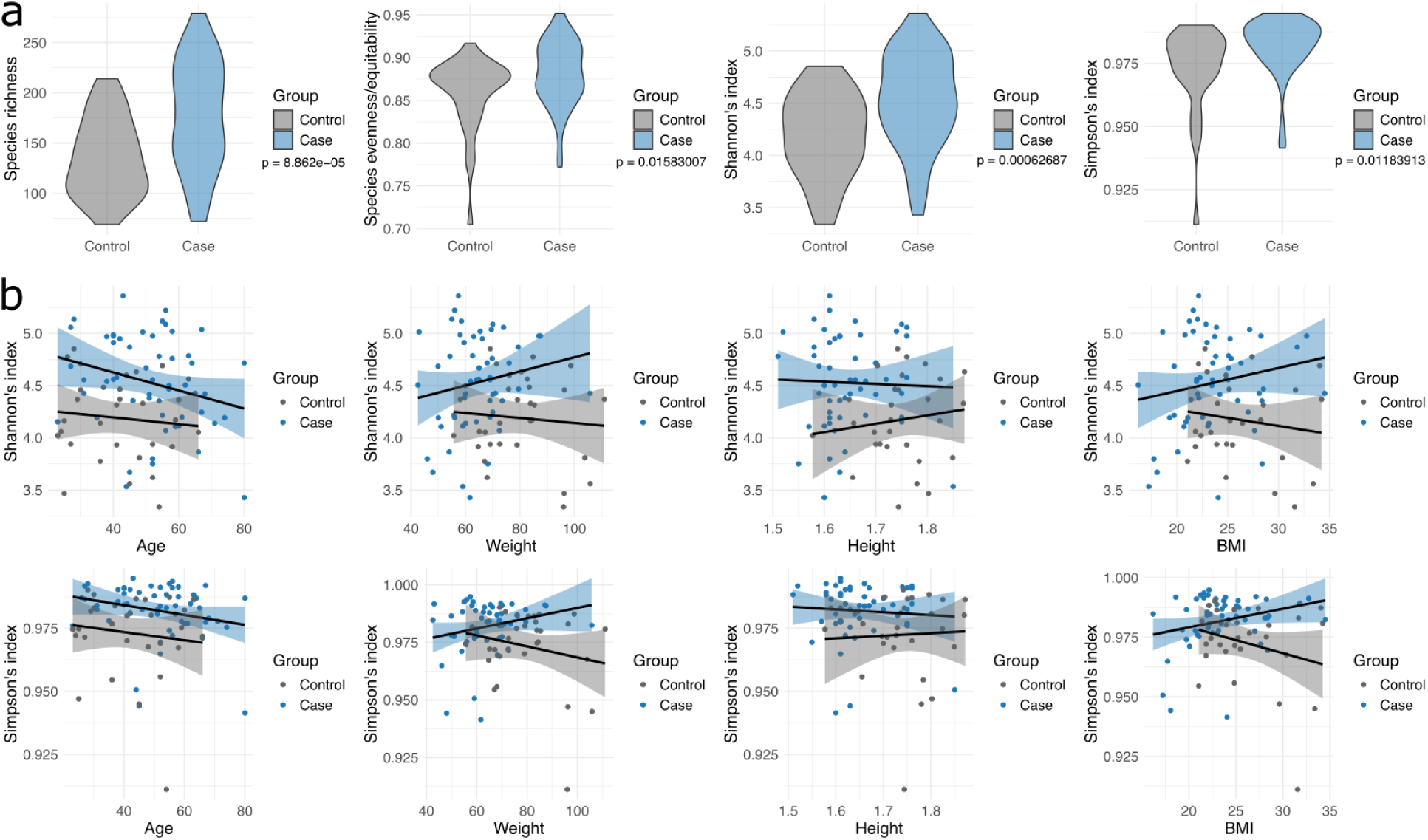
Alpha diversity. (a) Microbial diversity indices of CHA (blue) vs housed (gray) individuals. Indices include species richness, evenness (Shannon/log richness), Shannon, and Simpson. Diversity was significantly higher in CHA individuals across all indices (two-sided t-tests, *p* < 0.05). (b) Relationship between Shannon and Simpson indices and host covariates (age, weight, height, and BMI). In the CHA group, diversity increased with higher weight and BMI, while in controls, the trend was the opposite. Shaded areas represent 95% confidence intervals from linear models fitted separately to each group.

**Figure 2.**
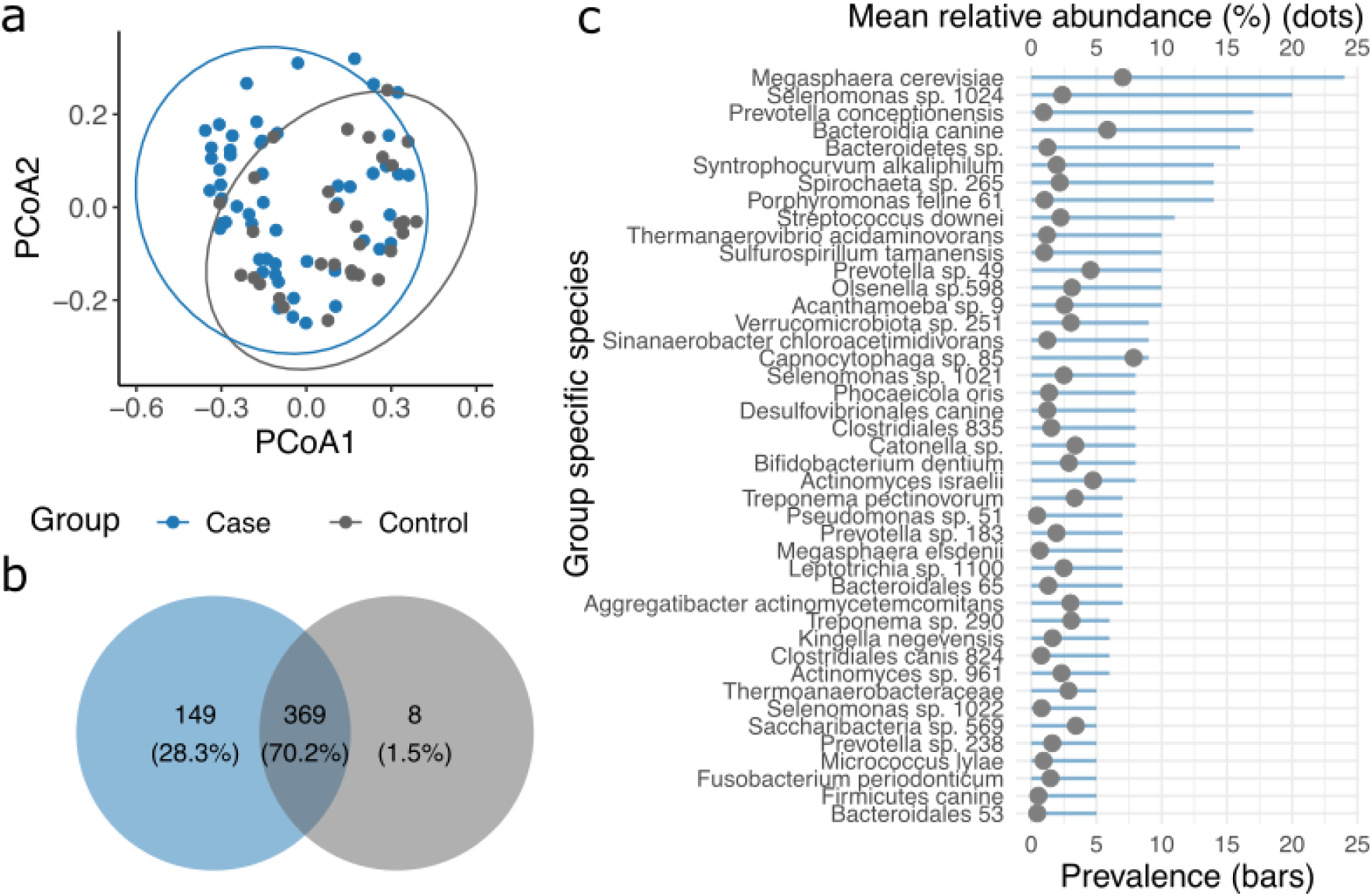
Differences in species composition between CHA and housed individuals. (a) Principal Coordinates Analysis (PCoA) based on Bray-Curtis dissimilarities using normalized abundances of all detected species. Each point represents one individual, colored by group (Case = blue, Control = gray). Confidence ellipses represent 95% intervals calculated independently for each group. (b) Venn diagram showing the number of unique and shared species between groups. (c) Prevalence and mean normalized relative abundance of CHA-specific species present in at least five individuals (prevalence ≥ 5). Normalized abundance is displayed as a percentage means per group. Blue bars represent species prevalence; gray dots indicate mean relative abundance.

The analysis of the differential abundance of the 369 species shared by the two groups is described in Figure 3. A volcano analysis distinguished those species significantly more abundant in either group (Figure 3a). The number of significantly more abundant species was markedly higher in cases than in controls, and within the top 10 more abundant, six were of the genus *Prevotella* (*Prevotella veroralis*, *Prevotella histicola*, *Prevotella falseni*, *Prevotella multisaccharivorax*, *Prevotella fusca,* and *Prevotella koreensis*). Whereas *Streptococcus sanguinis*, *Neisseria bacilliformis,* and *Actinomyces gerencseriae* were among the few species abundant in controls. In addition, the differential abundance of the 20 more abundant species in both groups was determined (Figure 3b), and *Capnocytophaga leadbetteri*, *Gemella haemolysans*, *Kingella denitrificans*, *Porphyromonas gingivicanis*, *Prevotella copri*, *Rothia mucilaginosa*, *Selenomonas flueggei* and *Streptococcus parasanguinis* were notably higher in cases, suggesting they are not only resilient to the harsh environment in CHA mouth but may even grow better. We also observed that *Actinomyces odontolytica*, *Capnocytophaga ochracea*, *Fusobacterium polymorphum*, *Granulicatella adiacens*, *Isoptericola chiayiensis,* and *Prevotella histicola* were notably lower, probably because they do not adapt efficiently, although they may still survive in this unique niche. In contrast, *Staphylococcus aureus* and *Fusobacterium hwasookii* were abundant in controls but absent in cases suggesting they are unable to survive in the CHA mouth.

**Figure 3.**
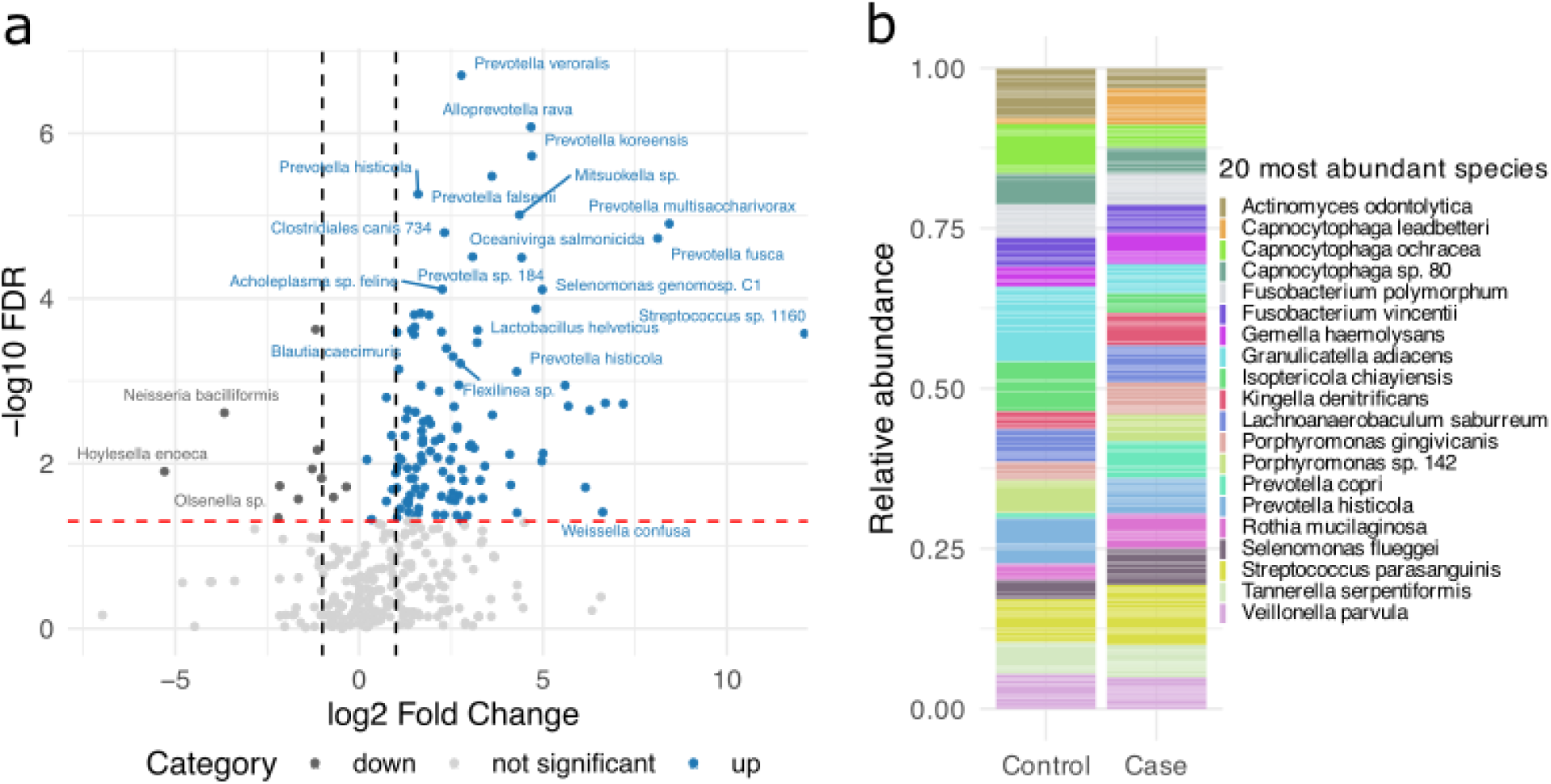
Differentially abundant species between CHA and housed individuals. (a) Volcano plot comparing the relative abundance of shared species between groups. Species significantly different are shown with adjusted p-values < 0.05 (red line) and a fold change threshold of |log₂FC| ≥ 1 (black lines). Only extremely enriched species in CHA individuals (blue) and controls (gray) are highlighted. (b) Relative abundance of the 20 most abundant species across all samples for cases and controls. Only species present in ≥ 2 individuals were included. Bars represent aggregated mean relative abundance per group; species are distinctly colored.

Furthermore, informative species were also determined with ML using Random Forest and LDA (Figure 4), and results with RF were similar to those with Volcano test, with *Prevotella veroralis*, *Prevotella histicola, Alloprevotella rava*, *Streptococcus sanguini*, *Prevotella korensis*, *Abiotrophia defectiva* and *Prevotella falseni* among the most informative (Figure 4a). On the other hand, with LDA *Leptotrichia massiliensis*, *Prevotella falsenii*, *Abiotrophia defectiva*, and *Megasphaera cerevisiae*, were the more informative for cases, whereas *Veillonella massiliensis, Actinomyces gerencseriae and Gemella haemolysans* were for controls (Figure 4b). A PCoA analysis using exclusively the informative species resulted in a clear separation of the groups (Figure 4c).

**Figure 4.**
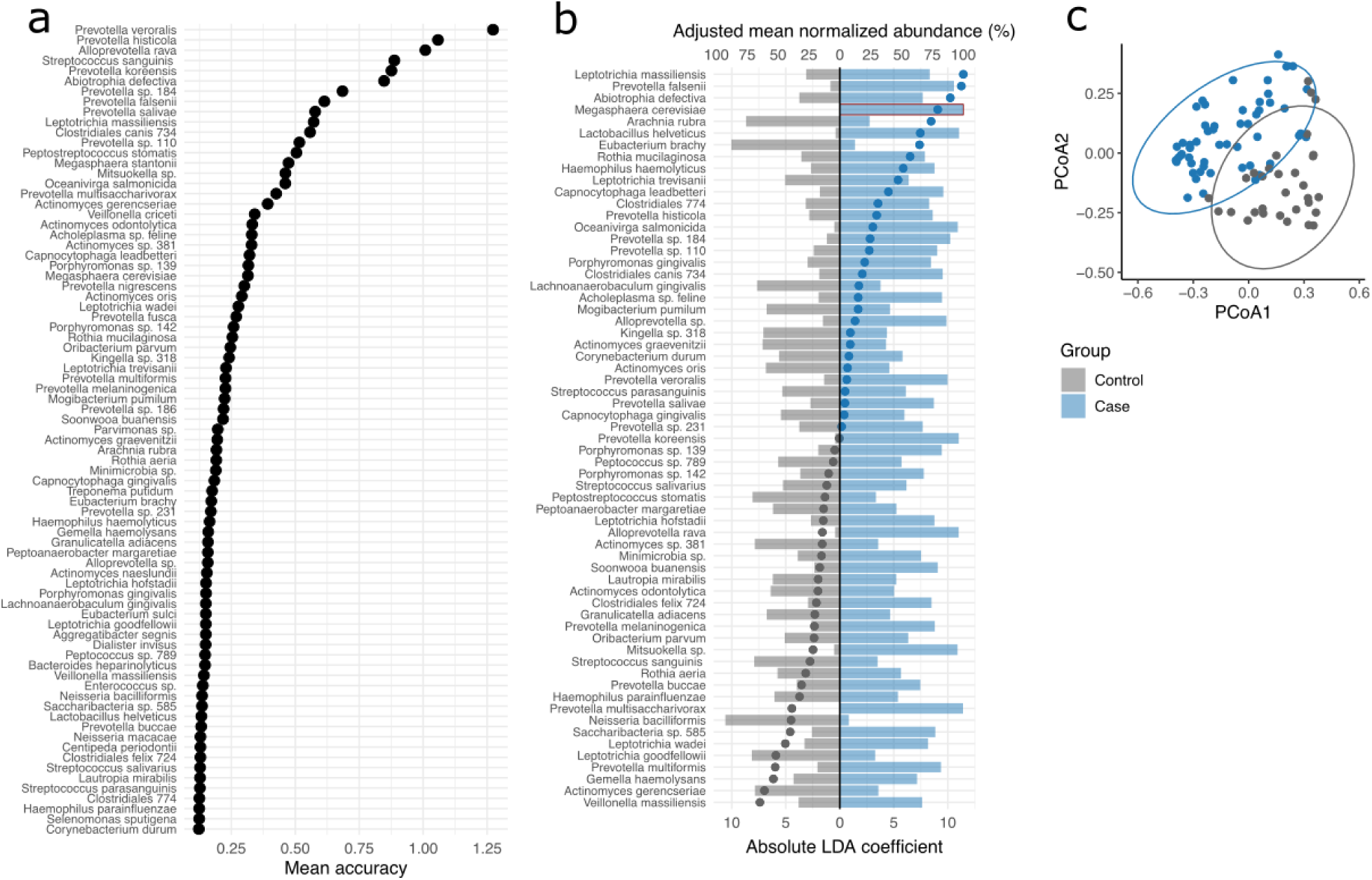
Identification of informative species. (a) Species importance from a 10-fold cross-validation Random Forest (RF) model. Informative species were defined as those above the 85^th^ percentile of mean accuracy (cutoff = 0.1227), yielding a model accuracy of 87%. (b) Linear Discriminant Analysis (LDA) of informative species selected by RF. Highly correlated species (Pearson |r| > 0.75) were removed to avoid multicollinearity. Bar lengths represent adjusted mean normalized abundance (%) per group; dots indicate absolute LDA coefficients. Group-specific species present in ≥5 individuals are marked with a red outline. (c) Bray-Curtis PCoA calculated from species selected by LDA. Ellipses represent 95% confidence intervals per group.

### Structure of bacterial communities

The interaction between the members of the bacterial community in each group was studied with network analyses (Figure 5). In the control group (Figure 5a) two main subclusters are observed, (red and blue) with strong links (thicklines) within subclusters and numerous links (thin lines) intra- and inter-subclusters. These subclusters include important nodes to maintain the net structure, *Corynebacterium durum*, *Prevotella buccae,* and *Abiotrophia defectiva*. Then there is a smaller subcluster (purple) where members show mostly negative links (red lines) with members of the other two subclusters, and includes *Prevotella histicola, Megasphaera stantonii* and *Rothia mucilaginosa*. These negative correlations are necessary to regulate the structural population of the oral microbiota and are important to maintain stable the bacterial communities. Furthermore, the thickness of several green lines in all subclusters suggests a strong dependence between the two species and a mark of a long-term relationship observed in mature stable bacterial communities.

**Figure 5.**
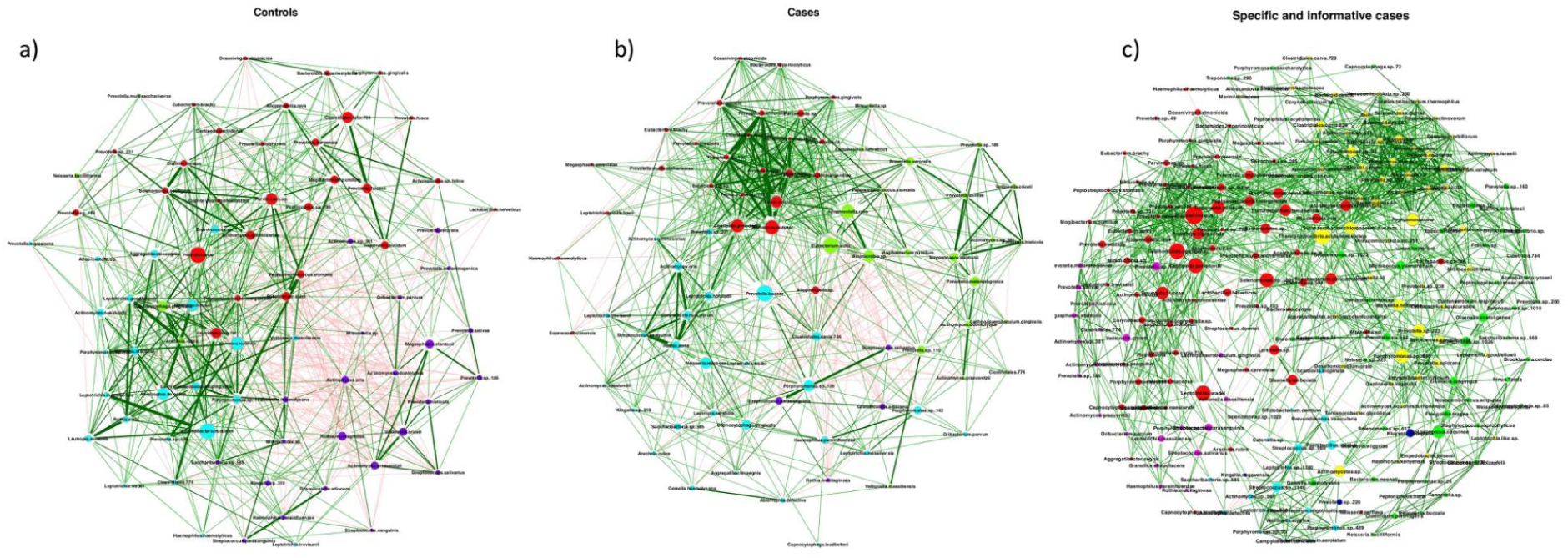
Abundance networks for housed (controls) and CHA (cases). (a) abundance network of controls, (b) abundance network of cases, and c) abundance network of case-specific species plus the informative species of cases obtained from Random Forest and LDA analysis. The networks were analyzed with Pearson method in the NetCoMi package. The size of the nodes was adjusted by betweenness, and a color was assigned to each subcluster within the network. Positive associations are shown with green connections and negative with red connections.

In the cases network (Figure 5b) four subclusters were apparent. The red cluster was densely connected with thick positive links with members of its own subcluster, but also with members of the other subclusters, and included *Selenomonas sputigena*, *Dialister invisus,* and *Centipeda periodontii* important for the net structure. *Megasphaera cerevisiae is a species* present exclusively in this CHA group (the most prevalent and abundant species). A second subcluster (green) contains three of the most informative species distinguishing cases from controls (*Prevotella veroralis*, *Prevotella histicola* and *Alloprevotella rava*) and *Eubacterium sulci*, important for the network structure. A third subcluster (blue) includes *Prevotella buccae*, one of the most important species for the stability of the network and one with the highest number of links (see below); *Clostridiales canis* a species present only in cases is also in this subcluster. Then a fourth subcluster (purple) where members present many negative links with species of the other subclusters, particularly *Streptococcus salivaris*, *Granulicatella adiacens,* and *Rothia mucilaginosa*. This last species also displayed negative links in the control group, whereas the other two also but to a lesser extent.

Finally, we aimed to understand how environmental, non-human-associated bacteria interacted with members of the oral microbiota. To explore this, we analyzed the network formed by the 149 case-specific species plus the 78 informative species present in cases (Figure 5c). Two main subclusters were apparent (red and yellow), containing most of the case-specific species and interacting with a dense net of links intra and inter-subclusters. These two subclusters also contained the species more important to sustain the network, including *Selenomonas sputigena*, *Dialister invisus*, *Sinanaerobacter chloroacetimidivorans,* and *Thermanaerovibrio acidaminovorans*. A third subcluster (green) with lower number of members, intermingles with the yellow and includes *Eikenella longinqua*, *Olsenella scatoligenes,* and *Micrococcus yunnanensis*. It also is important to note the high number of links that the case-specific species form with species present in both cases and asymptomatic controls, but with a low correlation (thin lines).

Notably, the prevalence of negative links differed between groups: in controls there were 0.29 negative links per one positive, in cases 0.12, and in the net of case-specific species 0.04, suggesting an important loss of regulatory interactions. Also, it is interesting to note that strong links (thick lines) are more frequent in controls than in cases and are uncommon in the case-specific network. These robust interactions in healthy microbiota are likely the result of long-term coevolution and ecological stability.

Another key finding is the difference in the number and type of nodes: while the control network contained 1,449 nodes, the case network had a higher count, with 1,531 nodes. Furthermore, it is also relevant with whom these nodes interact (Figure 6). For instance, *Alloprevotella rava* presented 13 nodes with the same species in controls and cases, but 32 exclusively in cases and only 1 specific for controls. In contrast, *Abiotrophia defectiva* presented 21 nodes exclusive for controls but only 12 for cases. *Prevotella buccae* presented 22 nodes, which share species in controls and cases, but 27 exclusively in cases and only 8 in controls (Figure 6).

**Figure 6.**
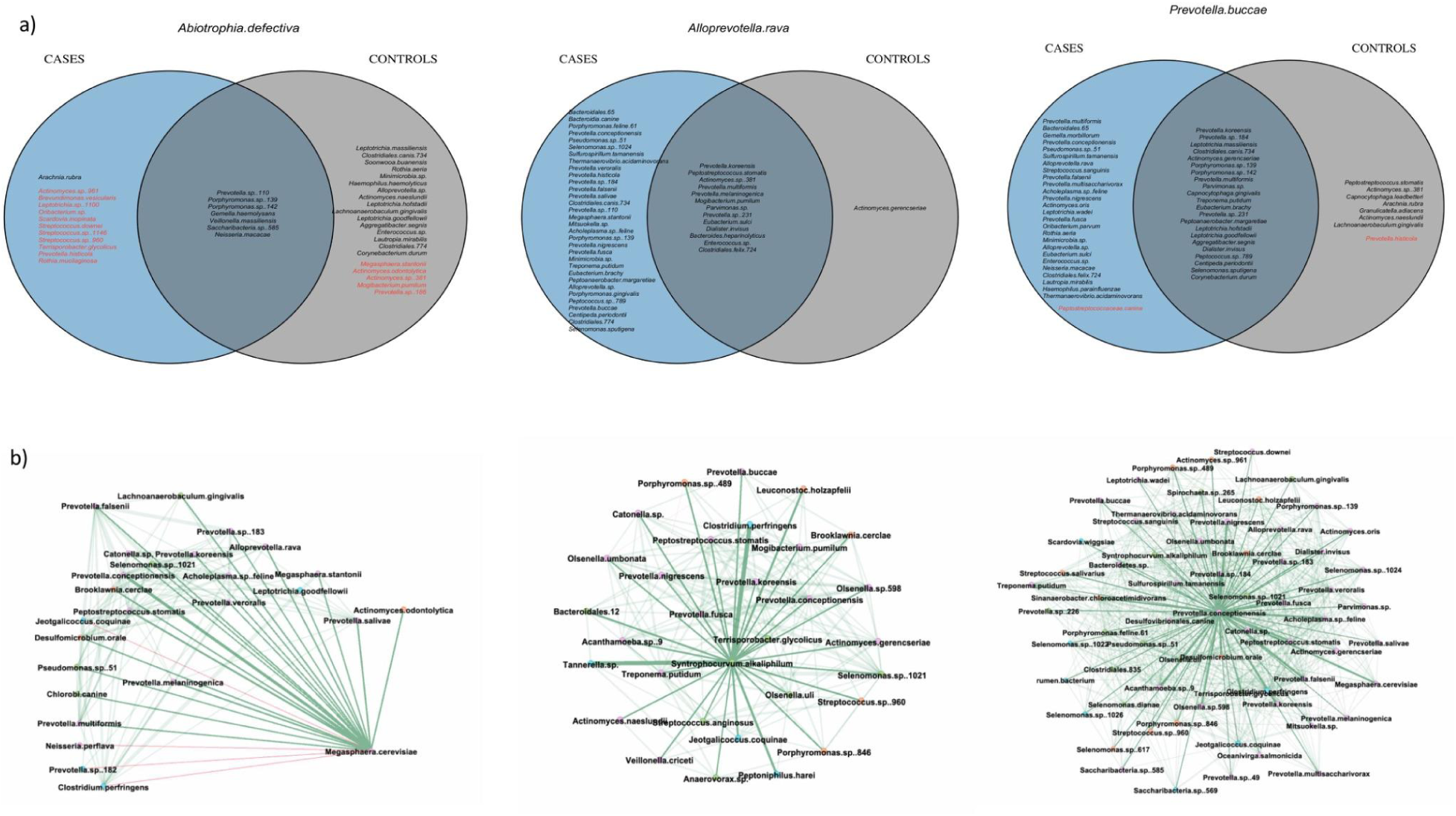
Representative diagrams with the number and type of links of some significant species in the networks. (a) The Venn diagram shows with whom the nodes *Abiotrophia defectiva, Alloprevotella rava*, and *Prevotella buccae* are interacting; in red are the negative links, while in black are the positive ones. (b) The networks display the links of *Megasphaera cerevisiae*, *Syntrophocurvum alkaliphilum*, and *Prevotella conceptionensis*, three species specific to CHA. In red are the negative links, in green the positive links.

Although not with strong links, the environment-extremophile species formed links with species of the human oral microbiota, as illustrated in Figure 6b.

### Estimation of the metabolic activity of the bacterial communities present in asymptomatic adults and in CHA

Figure 7 presents the projected functional profiles of bacterial communities from the oral cavity of individuals experiencing homelessness. A heatmap comparing the differential abundance of metabolic pathways between the CHA and control groups showed a significant enrichment in the CHA group for pathways related to the metabolism of fatty acids, sulfur, methionine and cysteine, terpenoids, carotenoids, and butyrate, as well as carbon fixation and homologous recombination (Figure 7a). In contrast, the “Control” group exhibits a predominance of pathways associated with signaling and transport processes, including quorum sensing, ABC transporters, and D-alanine metabolism. Differentially enriched metabolic pathways were also estimated with LDA with results very similar to the heatmap analyses (Figure 7b).

**Figure 7.**
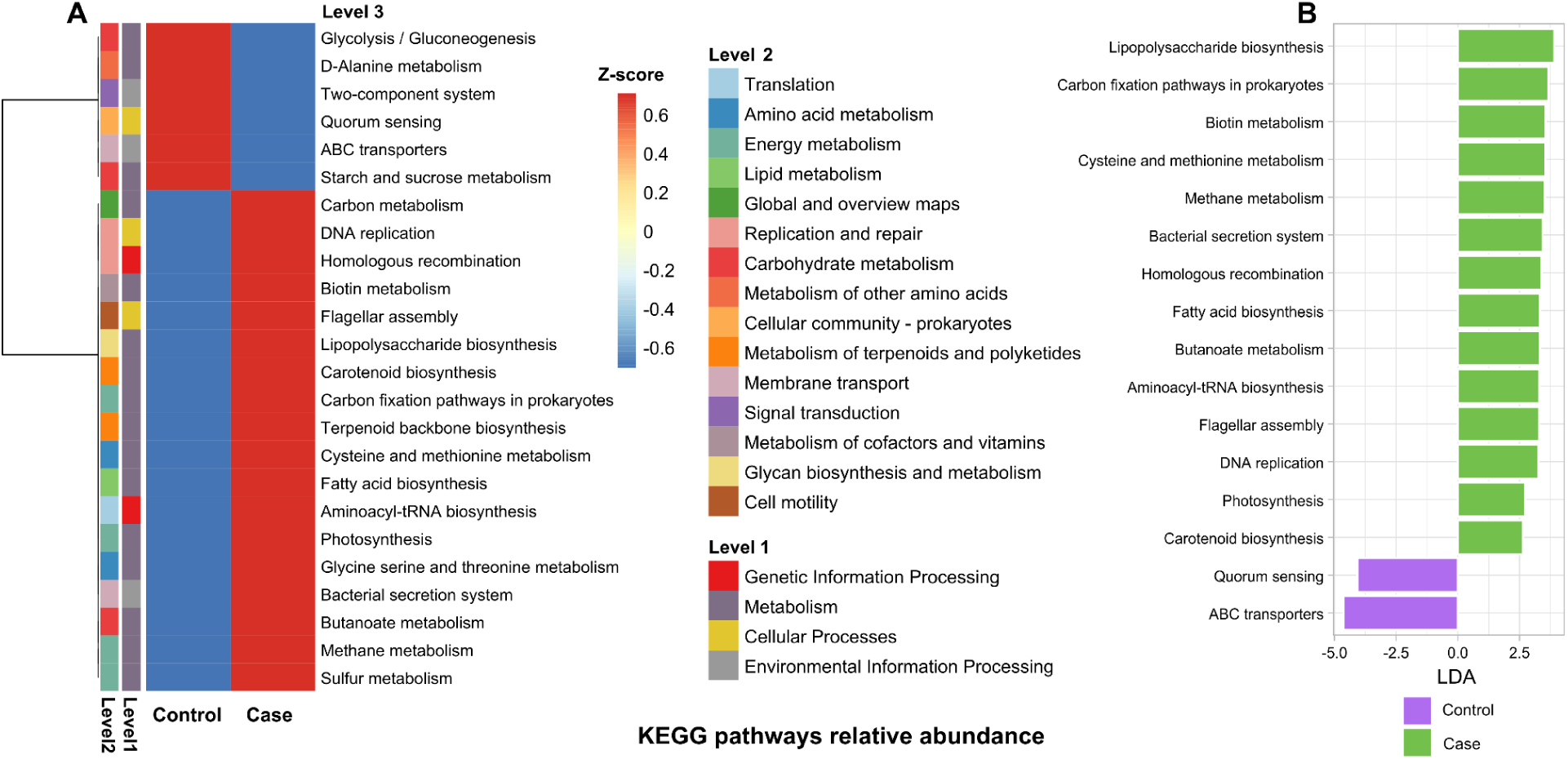
Metabolic activity of the bacterial community present in each group as inferred by Tax4fun2 software and KEGG database. (A) shows a heatmap with z-scores of the relative abundance of the metabolic pathways in level 1, 2, or 3 as defined in KEGG. (B) differentially abundant pathways between groups. Pathways enriched in the case group are indicated with a positive LDA score (mint green), and pathways enriched in the control group are indicated with a negative score. Only pathways with a significant LDA threshold of 2 and *p*-values < 0.05 are shown.

## Discussion

People in a homeless condition have more severe decay and missing teeth than the general population.It is estimated that around 83% of homeless had not had a dental cleaning in the previous 4 years (*website*, n.d. -c) and they are about 12 times more likely than individuals with stable housing to have dental problems, such as oral infections or edentulism (Lione et al., 2024) (Bernardino et al., 2021; de Pereira et al., 2014) (Bernardino et al., 2021) (Coles et al., 2011) (Wojcik et al., 2011). People living in a homeless condition continually expose their mouth to low-cost drugs like thinner - a mixture of oil-derived volatile organic compounds like toluene, xylene, acetone, and alcohols, all extremely harmful to human tissue. In addition, lack of oral hygiene results in a years-long chronic accumulation of toxic compounds in the oral cavity, leading to an extremely modified niche for bacteria. In the present study, we wonder what kind of bacteria may colonize such a harsh and toxic environment, and for that, we characterized the microbiota present in the saliva of a group of CHA individuals, aiming to understand the composition and networking of these bacterial communities.

Our work documents major differences between groups in all analyses tested. Richness, evenness, and diversity indexes were all higher in the CHA group an unexpected result indicative that the homeless-oral niche is not as restrictive for bacteria as we thought. Furthermore, we found many species (149) present exclusively in the CHA group, of which several of them were not previously reported in humans but in extreme environmental sources. In fact, as stated above, annotation using human microbiome databases left many ASVs unassigned.

The following examples illustrate the kind of bacteria colonizing the mouth of CHA people. *Megasphaera cerevisiae* was one of the most abundant species (around 6% of relative abundance) and the most prevalent (in 24 individuals), which suggests CHA got it from a common source. We learned that they frequently drink the same brand of sugarcane distilled with 20% ethanol and a mixture of volatile compounds, which probably contain alcohol resistant bacteria. *M. cerevisiae*, a strictly anaerobic species, is known for its ability to grow in beer, a very restricted media (Sakamoto & Konings, 2003; Santos, 2020) where low pH, ethanol content, limited carbon sources, hop bitter compounds, and high carbon dioxide are conditions that limit the growth of bacteria. *M. cerevisiae* has genes coding for multidrug efflux pumps and multiple copies of alcohol dehydrogenase that would explain its tolerance to high concentrations of ethanol (Bergsveinson et al., 2017).

Other identified extremophile anaerobic bacteria included *Syntrophocurvum alkaliphilum,* identified in a hypersaline soda lake in Siberia, is able to grow in high salt concentration and pH, capable to convert alcohols to methane and of anaerobic butyrate oxidation (Sorokin et al., 2016). *Thermanaerovibrio acidaminovorans* isolated from a granular methanogenic sludge, can ferment several amino acids to acetate and propionate and to grow lithoheterotrophically with H_2_ as electron donor and S^0^ as acceptor (Chovatia et al., 2009). *Sulfurospirillum tamanensis* isolated from a terrestrial mud volcano in Russia is an alkaliphilic bacterium that tolerates NaCl concentration of up to 14% and can use organic and inorganic compounds as electron donors or acceptors, it can even use arsenate for anaerobic respiration (Frolova et al., 2023). Of note, the three species described above were prevalent in 10 or more individuals. Furthermore, *Sinanaerobacter chloroacetimidivorans* isolated from an anaerobic acetochlor degrading reactor is capable of degrading the herbicide acetochlor and butachlor, with butyrate and acetate as the main fermentation products (Ventura et al., 2009a). *Bifidobacterium dentium* is tolerant to acid and resistant to chlorhexidine (Ventura et al., 2009a). These last two species were identified in 9 and 8 individuals, respectively. Thus, the above six extremophiles are species highly prevalent in the mouth of CHA individuals, tolerant to ethanol, salt, pH, acetochlor or chlorhexidine, properties that together could cooperate to thrive in the harsh environment of the homeless oral cavity. In fact, some of these case-specific species built a community-net within themselves but also with other species common in the human oral microbiome (see below) (Chovatia et al., 2009). All this seems to be an outstanding example of the plasticity of bacteria to form multimicrobial communities able to colonize hostile microenvironments that are rich in highly toxic compounds.

The other group of case-specific species included bacteria rare in humans but reported associated with disease and some also extremophiles. Thus, *Megasphaera elsdenii* is tolerant to high concentrations of salt, able to use indigestible substrates, and associated with type 2 diabetes and osteoporosis (Gao et al., 2024a) (Gao et al., 2024b; Ventura et al., 2009b), and *Phocaeicola oris* tolerant to acid and isolated from a case of oral squamous cell carcinoma (Chen et al., 2023). In addition, *Prevotella conceptionensis* was highly prevalent (present in 17 CHA) and is an anaerobe identified in systemic and tissular human infections (La Scola et al., 2011) (Toprak et al., 2019), *Streptococcus downei*, *Actinomyces israelii*, *Treponema pectinovorum*, and *Abiotrophia defective* have been reported associated with periodontitis and other tissular and systemic infections (Toprak et al., 2019) (Salman et al., 2019) (Pinilla et al., 2006) (Gholizadeh et al., 2017) (Sasaki et al., 2020). *Aggregatibacter actinomycetemcomitans* has been associated with aggressive periodontitis and tooth loss in adolescents (Fine et al., 2019). The variety of disease-associated bacteria present in the saliva of CHA people is outstanding, and the question is what the consequences for these individuals are.

CHA face severe oral health challenges, one of the most prevalent is dental caries (Kaste & Bolden, 1995) (de Pereira et al., 2014). Gingivitis and periodontitis are widespread (Bernardino et al., 2021; Freitas et al., 2019). Tooth loss is another major concern, with total edentulism, dental fractures, and the presence of numerous root remnants frequently observed. These conditions stem from untreated dental problems, but also from exposure to violence and accidents (Coles et al., 2011). Furthermore, oral infections and acute inflammation are common, often leading to dental abscesses. The retention of root residues, and underlying comorbidities—including hypertension, diabetes, psychological disorders, osteoporosis, and HIV—exacerbate these issues (Lione et al., 2024). Moreover, CHA individuals face an increased risk of developing oral cancer due to the frequent consumption of tobacco, alcohol, and other substances (Asgary, 2018). Collectively, these factors highlight the urgent need for targeted interventions to improve the oral health and overall well-being of this vulnerable population.

A key element in microbiota is the structure and networking of the bacterial communities, an issue poorly studied. We analyzed different properties of these bacterial communities, and correlation analysis showed marked differences between the two groups, including the species important to sustain the net, the nodes, edges, and subclusters. Also, some species differed drastically in the bacteria with whom they interacted in each of the groups, resulting in communities largely different in composition and structure. This highly dynamic behavior is driven in part by the nutrients, energy source, metabolic interactions, or production of antibiotics by members of the bacterial communities. Another important factor is the composition of the microenvironment in the mouth, which is strongly influenced by diet and habits that are drastically different in CHA people, resulting in extremely toxic conditions and unfavorable for the usual human oral microbiota. Thus, it was expected to identify uncommon bacteria in the saliva of these people, although not that many (132 SVUs case-specific vs 6 control-specific) and not that extremophilic (tolerant to alcohol, salt, acid, herbicides or oil solvents).

The presence of environmental extremophilic bacteria in the oral cavity of homeless individuals reveals a remarkable ability of the microbiome to adapt to a human hostile environment (Douglas et al., 2014). In this context, some metabolic pathways play a key role in the successful adaptation of these microorganisms. Increased activity of some metabolic pathways was observed in the case group. The glycolysis/gluconeogenesis pathway may represent a mechanism for metabolic self-organization under extreme fluctuations in nutrient availability. It allows for the adjustment of central carbon fluxes when energy is limited, avoids futile cycling, and ensures tight regulation of central metabolism—an essential feature for survival (Schink et al., 2022). D-alanine metabolism has been associated with adaptation to extreme environments (Yu et al., 2022). Quorum sensing plays a key role in microbial communication under such conditions (Kaur et al., 2019). ABC transporters and two-component systems are also involved in metabolic adaptation to harsh environments (Shulami et al., 2007). Additionally, starch and sucrose metabolism can enhance the formation of adherent biofilms, promote the production of water-insoluble polysaccharides, and contribute to environmental acidification (Duarte et al., 2008).

Among the pathways enriched in homeless microbiota, carbon fixation is important for autotrophs adapting to diverse environments. Thus, CO₂ assimilation might be an option for a carbon source in a nutrient-limited environment (Lemaire et al., 2020). Lipopolysaccharide biosynthesis is essential for the structural integrity of Gram-negative bacteria (Sperandeo et al., 2019), providing stability in extreme environments. Other metabolic pathways, such as fatty acid biosynthesis enable membrane adaptation to osmotic pressure or temperature changes (Suutari & Laakso, 1994).

This study has certain limitations, including a relatively small sample size. Individuals experiencing chronic homelessness often form fluid and unstable communities that are difficult to track and engage in long-term studies. This is partly due to health vulnerabilities, limited access to services, and the social exclusion they regularly face. Despite these challenges, the sample size was sufficient to detect clear and significant differences in microbiome composition between groups. Moreover, the consistent presence of specific extremophilic taxa - such as *Megasphaera cerevisiae* - supports the robustness of our findings.

CHA is a population that is constantly increasing worldwide as a result of the evolution of our societies and of the exponential growth of migrant communities that end up living in homeless conditions in foreign countries where they are seen and treated as social outliers. Whereas the Organization for Economic Co-operation and Development (OECD) estimated that at least 2 million people in OECD countries were in a homelessness situation in 2023 (*Website*, n.d.-b), projections estimate that people in a homeless condition may increase nearly triple by 2030 (Santos, 2020). No doubt the world must pay attention to CHA people and act to rescue them from the dark side of society.

In summary, our results document the outstanding adaptation of environmental extremophile bacteria to colonize hostile human niches continuously exposed to toxic compounds. It is highly probable that these bacterial communities have evolved to successfully adapt to this years-chronic scenario where extremophile-environmental and human bacteria now share a very unusual niche product of the evolution of modern societies.

